# An Experimental and Multiphysics Simulations Study of *Clostridium carboxidivorans* sp. 624 for Acid and Alcohol Production in CO_2_/H_2_

**DOI:** 10.1101/2025.02.24.639814

**Authors:** Yong Wei Tiong, Ashriel Yong, Anjaiah Nalaparaju, Hoang-Huy Nguyen, Li Quan Koh, Roong Jien Wong, Ron Tau Yee Lim, Dave Siak-Wei Ow, Jia Zhang, Van Bo Nguyen, Fong Tian Wong, Yee Hwee Lim

## Abstract

Biotechnological advances in CO_2_ utilization through gas fermentation offer a sustainable alternative to energy-intensive chemical processes. This study investigates and optimizes fermentation dynamics of *Clostridium carboxidivorans* sp. 624 for enhanced production of C_2_–C_6_ acids *via* combined experimental and computational methods. A systematic experimental evaluation of temperature and medium conditions, along with time-course analysis elucidates metabolic pathway regulation. Experimental optimisation, complemented by modelling, identified gas-liquid volumetric ratios V_L_/V_G_ = 4 as optimal for maximizing longer-chain acids production, attributed to enhanced gas solubility and substrate bioavailability. Additionally, a hybrid model combining dynamic-kinetic model and computational fluid dynamics (CFD) successfully predict product formation. By capturing spatial inhomogeneities in gas-liquid interactions, the model also provides critical insights for optimizing fermentation performance and establishes a framework for improving process design parameters for CO_2_/H_2_ fermentation, paving the way for scalable and efficient CO_2_-based bioprocesses.

## 1. Introduction

Since the Industrial Revolution, human activities have significantly raised atmospheric CO_2_ concentrations, primarily due to the burning of fossil fuels. Consequently, CO_2_ levels have increased from approximately 277 ppm in 1750 to 421 ppm in 2022, contributing to global climate change and its associated environmental impacts (Raganati and Ammendola, 2024). To mitigate emissions, the Paris Agreement targets a 50–100% reduction by 2030 and beyond (Dziejarski et al., 2023). Surrounded by this, biotechnological CO_2_ utilization is emerging as a sustainable alternative to energy-intensive chemical catalysis (Liew et al., 2016), leveraging microorganisms to convert CO_2_ into valuable bio-based products and advancing a circular economy (Ricci et al., 2021).

One promising approach is anaerobic CO_2_ gas fermentation, in which clostridial acetogens utilize the Wood-Ljungdahl pathway (WLP), also known as the reductive acetyl-CoA pathway, to fix CO_2_ (Acuña et al., 2024; Antonicelli et al., 2023; Ntagia et al., 2021). The resulting acetyl-CoA undergoes further metabolic reactions to produce organic acids and alcohols (Sun et al., 2019). This process occurs in two distinct phases: acidogenesis (organic acid production) and solventogenesis (conversion to alcohols), where CO_2_ serves as the electron acceptor and H_2_ as the donor, enabling CO_2_ fixation and sustainable bioproduction (Mariën et al., 2024). Despite advancements in syngas fermentation using CO, CO_2_, and H_2_ (Doll et al., 2018; Fernández-Naveira et al., 2019; Phillips et al., 2015), CO_2_/H_2_ fermentation remains in the early stages of bioengineering, primarily yielding acetate (Molitor et al., 2019) and low ethanol titers (Heffernan et al., 2020). Expanding production to higher-value medium-chain C_4_ and C_6_ bio-based chemicals is essential for improving the economic feasibility of CO_2_ fermentation (Thunuguntla et al., 2024).

Among acetogenic bacteria, *Clostridium carboxidivorans* sp. 624 is known for producing a wide range of C_2_ to C_6_ acids and alcohols, including acetate, butyrate, ethanol, butanol, hexanoate, and hexanol, when utilizing syngas or CO-based feedstocks (Oh et al., 2022; Phillips et al., 2015). In syngas fermentation (CO/H_2_/CO_2_ [70/20/10]), this strain produced mainly ethanol (∼3.00 g/L) and hexanol (0.94 g/L) (Phillips et al., 2015), while a different syngas mixture (CO/H_2_/CO_2_ [30/40/30]) yielded 4.32 g/L total alcohols, with hexanol (1.90 g/L) as the dominant product (Oh et al., 2022). However, research on pure CO_2_/H_2_ fermentation remains limited. A recent study employing Adaptive Laboratory Evolution (ALE) reported modest metabolite yields, including acetate (0.82 g/L), butyrate (0.38 g/L), and hexanoate (0.40 g/L), alongside low alcohol titers (0.23–0.27 g/L) under CO_2_/H_2_ fermentation (Antonicelli et al., 2023). These findings highlight challenges such as limited metabolite titers and variability in product distribution, underscoring the need for optimization in microbial pathways and fermentation parameters to enhance CO_2_-based biomanufacturing. Importantly, CO_2_/H_2_ fermentation could also offer a promising direct utilization pathway for CO_2_, without requirements of CO inputs.

In addition to experimental optimization, modelling and simulation can play a crucial role in optimizing bioprocesses by analyzing parameters *in silico*, minimizing trial-and-error experimentation, and accelerating process development. In C_1_ gas fermentation by the clostridial microbiome, dynamic-kinetic model captures microbial metabolic pathways and integrates mass balance to generate time-dependent metabolite profiles. Computational tools, including dynamic simulations and computational fluid dynamics (CFD), assess mass transfer, which is critical in gas-to-liquid fermentation systems (Puiman et al., 2022). Modelling CO_2_ and H_2_ spatial distribution provides insights into their impact on microbial growth and metabolite production (Singh et al., 2024). Despite advancements in gas fermentation modelling, models specifically tailored for CO_2_/H_2_ fermentation in clostridial strains producing C_2_-C_6_ metabolites remained unstudied, with most studies focused on CO as the primary substrate (Ruggiero et al., 2022; Younesi et al., 2005).

Herein, this study systematically evaluates CO_2_/H_2_-based fermentation of *C. carboxidivorans* sp. 624 towards the production of C_2_–C_6_ acids and alcohols through a combined experimental and *in silico* approach. A time-course study examines the metabolic dynamics of C_2_–C_6_ acid and alcohol production, providing insights into pathway regulation and intermediate accumulation. Using the experimental data, an integrated model featuring a hybrid CFD simulation-first principles kinetic model was developed to predict product formation. By capturing spatial inhomogeneities in gas-liquid interactions, the model also provides critical insights for optimizing fermentation performance and establishes a framework for improving process design parameters for CO_2_/H_2_ fermentation, paving the way for scalable and efficient CO_2_-based bioprocesses.

## 2. Materials and Methods

### 2.1 Microorganism and medium preparation

*C. carboxidivorans* sp. BAA-624 was purchased from American Type Culture Collection (ATCC, USA). The modified PETC medium was prepared by dissolving the following components per liter of deionized water: 0.10 g KCl, 0.20 g MgSO_4_·7H_2_O, 0.80 g NaCl, 0.10 g KH_2_PO_4_, 0.02 g CaCl_2_·2H_2_O, 1.00 g yeast extract, and 2.00 g NaHCO_3_. Additionally, 10 mL of modified Wolfe’s metals solution (Med 1788) and 10 mL of Wolfe’s vitamin solution (MD-VS; Med 1019) were added to supplement the medium. Reinforced Clostridial Medium (RCM, Thermo Scientific™ Oxoid™ Reinforced Clostridial Medium) was prepared from a dehydrated powder (Thermo Scientific™ Oxoid™, catalog number CM0149B) according to the manufacturer’s instructions.

Fermentation was conducted in 30 mL sterilized serum bottles, each containing 15 mL of PETC/RCM medium and 15 mL of headspace. The headspace was flushed for 1 minute with a CO_2_/H_2_ gas mixture (20/80), after which the pressure was equilibrated to 150 kPa (1.5 bar) with the same gas. To maintain consistent gas substrate availability, the headspace of each culture bottle was replenished daily by flushing with fresh CO_2_/H_2_ (20/80) for 1 minute and equilibrated to 150 kPa. Fermentation was carried out at 25°C, 30°C, and 37°C under mild agitation at 200 rpm using an orbital shaker (Yih Der, LM-570RD). Control cultures were prepared by sparging with N_2_ gas following the same procedure as the experimental conditions, which involved sparging for 1 minute and equilibrating to 150 kPa. N_2_ gas controls are presented in **Fig.1-2** and **Fig S1-S2**. All data are reported as mean values from triplicate experiments ± standard error of the mean.

### 2.2 Analytical methods

#### Biomass analysis

The samples from fermentation were analysed with a multimode microplate reader (Thermo Scientific™ Varioskan™ LUX, USA) for optical density (OD=800nm) measurements, and a correlation factor was used to obtain cell dry weight (CDW) concentrations. To obtain CDW conversion factor, a sample volume with known OD was pelletised and supernatant discarded. The pellet was dried at 60 °C for 48 hours followed by gravimetric analysis to determine CDW% per volume. Maximum specific growth rates were estimated by applying a non-linear regression to the experimentally obtained data for CDW concentrations.

#### Metabolites analysis

C_2_-C_6_ metabolites (ethanol, butanol, hexanol, acetate, butyrate, and hexanoate) from fermentation were analysed using a gas chromatography (GC) (Agilent 6890N GC, Agilent Technologies, USA) with a flame ionization detector (FID) and fused silica capillary column (DB-WAX 60 m length x 0.53 mm diameter x 1.00 µm film thickness). Helium was used as the carrier gas. The initial oven temperature was set at 35 °C with a holding time of 4 min, and then gradually increased to 220 °C at a rate of 10 °C/min, and held at the final temperature for 6 min. The injector and detector temperatures were both set to 300°C. Prior to GC analysis, the sample was centrifuged, and the supernatant was filter through a 0.22 μm filter. A 0.50 mL portion of the supernatant was then acidified with 0.02 mL of 0.60 N HCl before proceeding with the analysis. Production yields of the C_2_-C_6_ metabolites reported herein have been referenced to 0 at t = 0 hour.

### 2.3 Kinetic model of microbial reactions

The biochemical reactions involved in the conversion of CO_2_/H_2_ to C_2_-C_6_ acids and alcohols by *C. carboxidivorans* sp. 624 during both the acetogenic and solventogenic phases, via the WLP and reverse *β*-oxidation, are presented in equations below (**eq. 1-6**).

Acetogenic phase:

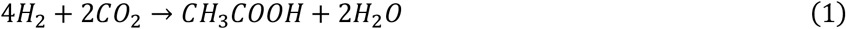

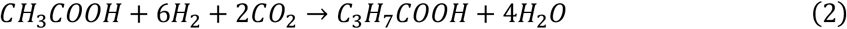

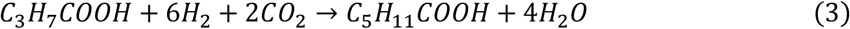

Solventogenic phase:

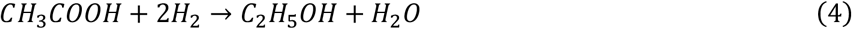

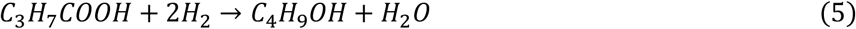

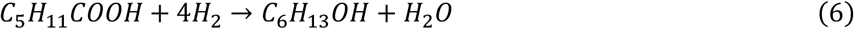

To describe the rates of acid production (q_AC_, q_BA_, and q_HA_), modified Monod equations that incorporate the inhibition effects of non-substrate fatty acids were applied (**eq. 7-9**). The conversion rate of acids in the presence of H_2_ to produce alcohols (q_E_, q_B_ and q_H_) were expressed as Monod kinetic for substrates (**eq. 10-12**).

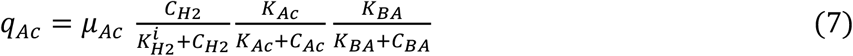

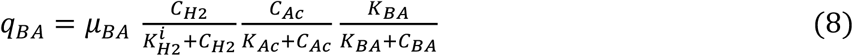

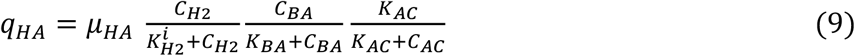

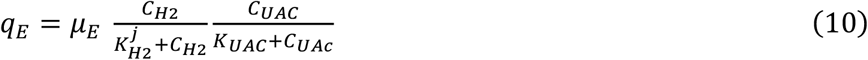

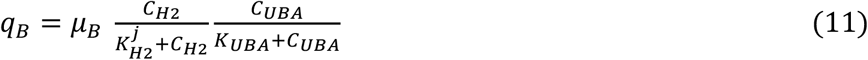

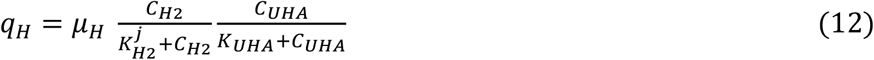

where µ_i_ (i=AC, BA, HA) are the maximum uptake rates of biomass for acids production, µ_j_ (j=E, B, H) are the maximum production rates of alcohols, 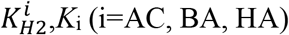 are affinity constants for biomass, 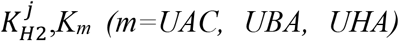 are saturation constants of H_2_, undissociated acids. *C*_n_ (n=AC, BA, HA, H2, UAC, UBA, UHA) concentration in liquid phase. (Abbreviation: AC-acetate, BA-butyrate, HA-hexanoate, E-ethanol, B-butanol, and H-hexanol)

The component mass balance equations for the gas and liquid phases are provided in **Table S1**. These first-principles 0D dynamic-kinetic model equations were numerically solved to generate time profiles for product formation, and kinetic model parameters were estimated to achieve optimal representation of the experimental data. The mass transfer coefficient (K_L_a) and Henry’s constant (K_H_) for H_2_ were obtained from literature sources (Manna et al., 2024).

### 2.4 CFD simulation for gas fermentation in serum bottle

To model multiphase flows in bioreactors, the Eulerian-Eulerian solver for multiple incompressible fluid phases in OpenFOAM (Weller et al., 1998) was employed for transient simulations. This solver treats each phase as a continuous fluid with distinct volume fractions, making it suitable for modelling and analysing complex fluid interactions in scenarios such as gas-liquid-cell flows and bubbly flows. Given the critical role of mass transfer in controlled aeration and microbial metabolism, the Higbie mass transfer model was integrated into the OpenFOAM framework, demonstrating superior accuracy compared to the existing spherical and Frössling gas-liquid mass transfer models (Mast et al., 2024). The detailed equations for this implementation were provided in **Eq. 13–16**.

In the current simulation framework, the biological model serves as a phenomenological representation of the complex metabolic processes of a specific microorganism. To enhance computational efficiency, the reaction model was simplified while retaining key kinetic phenomena necessary for quantitative experimental validation. This reduced-order model was then integrated with the multiphase flow model to capture essential biochemical interactions. The evolution of microbial and species concentrations follows the kinetic model detailed in **Table S1**. The system of ordinary differential equations governing these processes was solved using an implicit integration solver, while species transport was coupled with a second-order explicit Runge-Kutta scheme to ensure numerical stability and accuracy.

The multiphase CFD model is described as follows:

The mass conservation equation for each phase *i* is given by

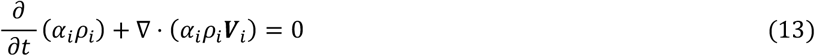

and the momentum equation for each phase i is given by

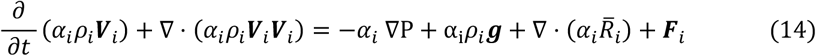

where *ρ*_*i*_ is the density, *V*_*i*_ is the phase velocity, P is the mean pressure, 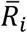 is the combined viscous and Reynolds stresses for phase i; and *F*_*i*_ is the interface momentum transfer term, which is the sum of all the interfacial forces (lift, drag, virtual mass, wall lubrication, turbulent-dispersion forces, etc) acting on phase I from the other phase. It is noted that this term will influence bubble rise velocity and has large effect on gas hold-up calculation.

The transport of chemical species is modelled with passive scalar in each phase ad is given by

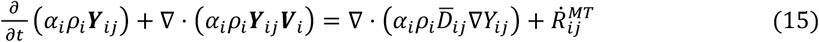

Here *Y*_*ij*_ represents the mass fraction of species j in phase i,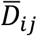 is the total species diffusion coefficient including both molecular and turbulent diffusion, and 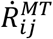 is the interface mass transfer term for species j in phase i.

The Higbie mass transfer equation is given by,

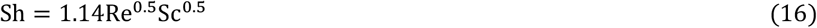

where Re is Reynold number, Sc is the Schmidt number which was used to characterize the mass diffusion and convection processes.

## 3. Results and Discussions

### 3.1 C_2_-C_6_ smetabolites production

The growth profiles of *C. carboxidivorans* sp. 624 in both PETC and RCM media at different cultivation temperatures of 25°C, 30°C, and 37°C, were investigated using either N_2_ or CO_2_/H_2_ (20/80) gas substrates at a headspace pressure of 150 kPa (RCM medium in **Fig. 1**, PETC medium in **Fig. S1**). The optimal temperatures for *C. carboxidivorans* have been reported differently in previous literature. Some studies suggest optimal growth occurs between 37°C and 40°C (Liou et al., 2005). Other research indicates that lower temperatures promote the production of acetate and ethanol, as well as facilitate carbon chain elongation (Ramió-Pujol et al., 2015). However, in our study, the *C. carboxidivorans* sp. 624 generally prefers an optimal temperature of 25°C. Comparison of the C_2_ metabolites production between N_2_ and CO_2_/H_2_ (20/80) gas substrates (RCM medium in **Fig. 1**, PETC medium in **Fig. S1**) suggests evidence of CO_2_ assimilation via WLP which corroborates with previous work (Antonicelli et al., 2023).

**Fig. 1.**
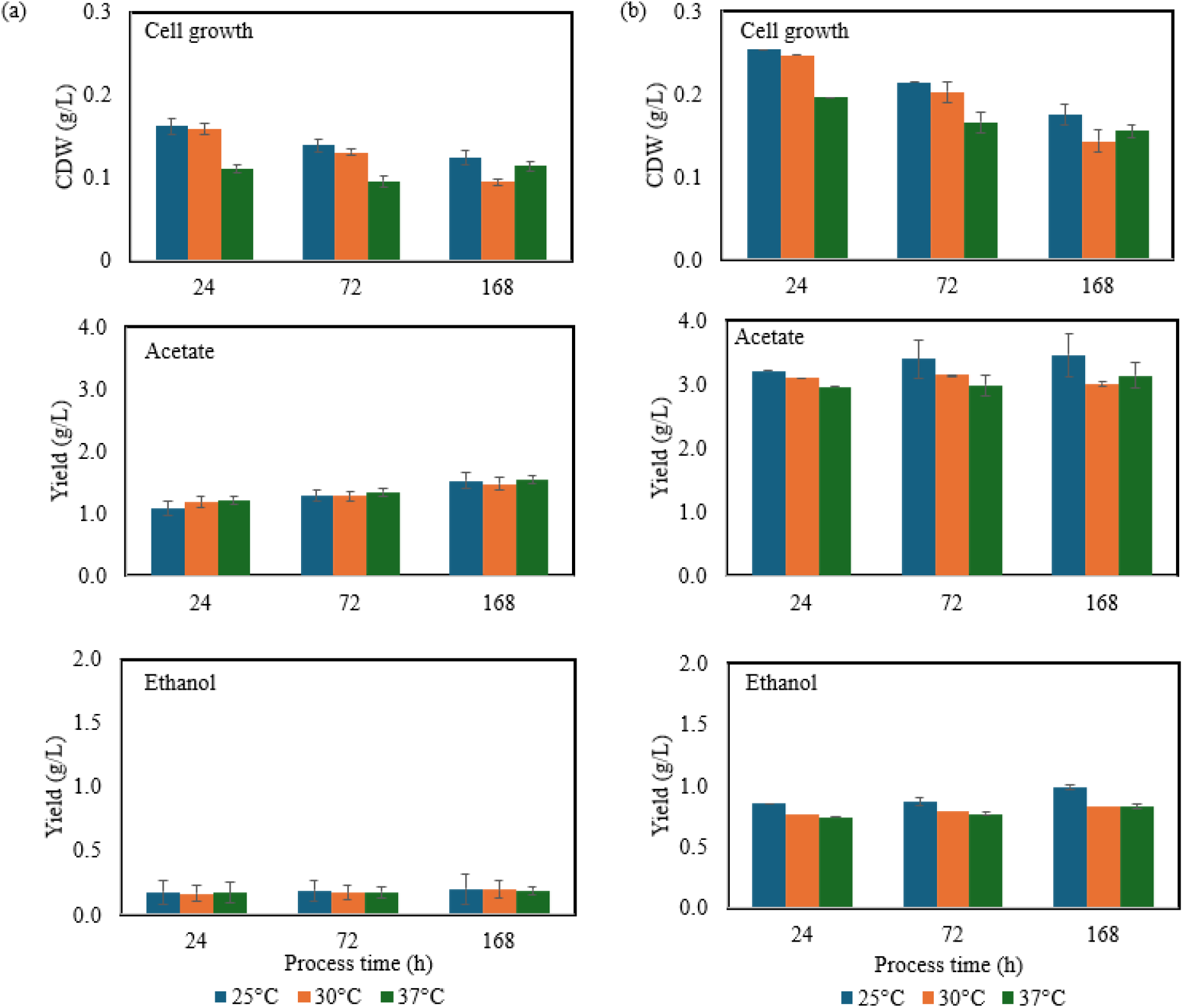
Cell growth and production of C_2_ metabolite analysis of *C. carboxidivorans* sp. 624 fermentation under **(a)** N_2_, and **(b)** CO_2_/H_2_ (20/80) in RCM media at 25°C, 30°C, and 37°C.

To gain a deeper understanding of the metabolite production, detailed sampling of C_2_-C_6_ metabolites was conducted at 24, 48, 72, and 120 hours, in RCM media at 25°C under CO_2_/H_2_ (20/80) atmosphere (**Fig. 2** and **Fig. S2**). The CDW results (**Fig. 2a**) indicated that cell growth peaked by 24 hours (exponential phase) before gradually declining and ultimately reaching the stationary phase at 120 hours. This trend suggests that the initial cell proliferation was primarily supported by organic nutrients in the media. Despite the decline in cell growth, continued metabolite accumulation beyond 24 hours implies sustained metabolic activity, driven by CO_2_ assimilation via the WLP. This observation is consistent with findings by Antonicelli et al. (2023), who reported early exponential growth followed by a delayed cell growth increase beyond 10 days.

**Fig. 2.**
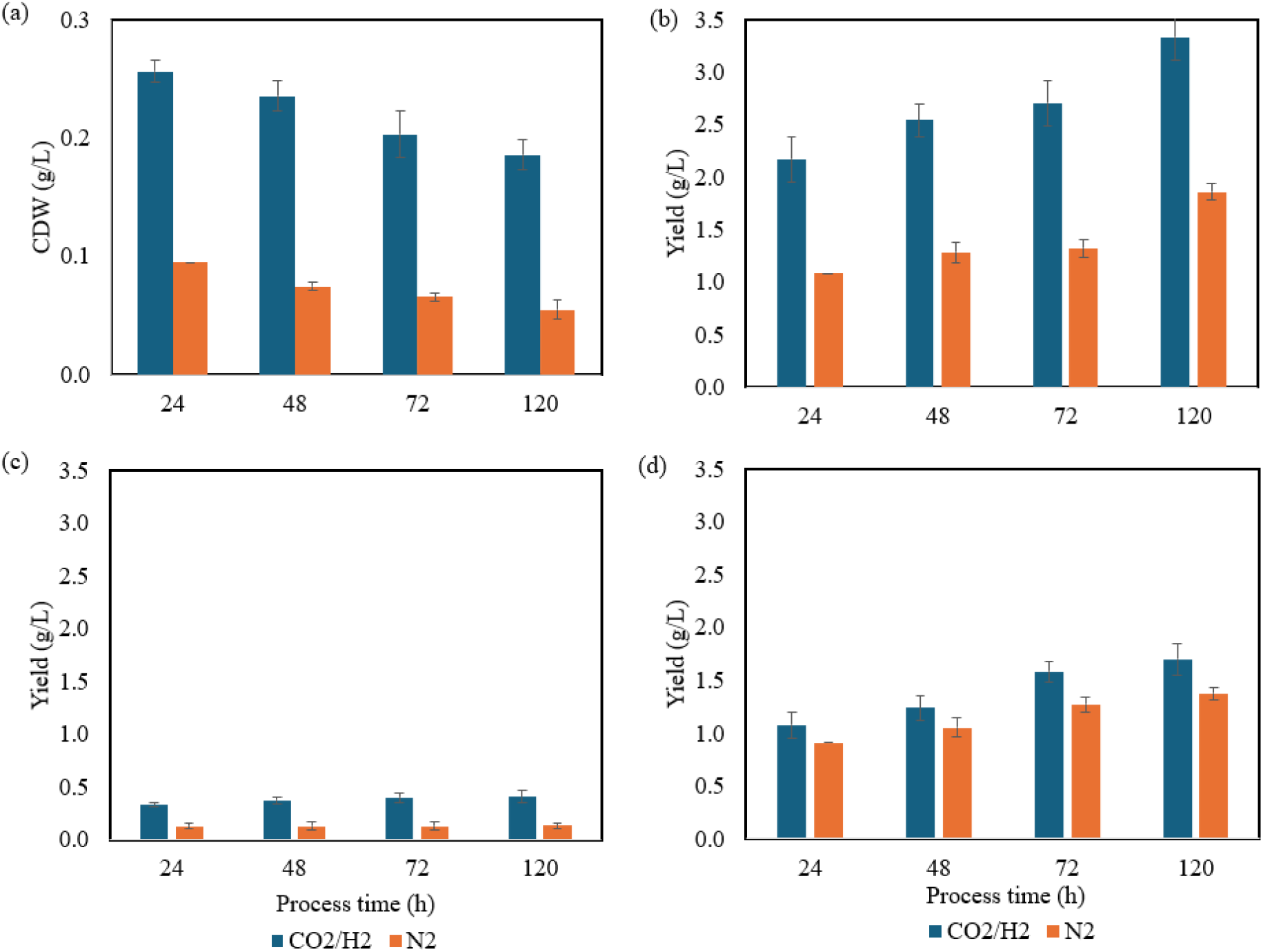
*C. carboxidivorans* sp. 624 fermentation with CO_2_/H_2_ (20/80) or N_2_ in RCM medium at 25°C: **(a)** Cell growth; Production yields of **(b)** Acetate, **(c)** Butyrate, and **(d)** Hexanoate.

Among the acid metabolites production, acetate was the primary product during the exponential growth phase, reaching a notable yield of 2.17±0.22 g/L in RCM by 24 hours (**Fig. 2b**). Acetate yield continued to gradually increase, reaching 3.33 ± 0.21 g/L in RCM by 120 hours under CO_2_/H_2_ conditions, likely driven by sustained metabolic activity and carbon fixation. The higher acetate production in the presence of CO_2_/H_2_, relative to the N_2_ control (1.08±0.09 g/L at 24 hours and 1.86±0.08 g/L at 120 hours, **Fig. 2b**) underscores the contribution of gas-phase CO_2_/H_2_ to acetate production. The slower rate of increase after 24 hours could be attributed to the inhibitory effects of accumulating byproducts, such as butanol, hexanoate, and hexanol (Oh et al., 2022). Similar trends for butyrate and hexanoate production were observed, where production was significant in the first 24 hours and slowed down after. Butyrate yield (**Fig. 2c**) remained relatively limited (<0.50 g/L) throughout the fermentation period, peaking at 0.41±0.06 g/L by 120 hours. Hexanoate production (**Fig. 2d**) followed similar trends to acetate, reaching an initial yield of 1.08±0.12 g/L within 24 hours and a final yield at 1.70±0.15 g/L by 120 hours. In terms of alcohols, their production remains minimal (<0.5 g/L) throughout the fermentation, indicating that solventogenesis pathway is not preferred under the investigated conditions (**Fig. S2**). Amongst which, ethanol is the most dominant solvent produced, compared butanol and hexanol (<0.1 g/L).

### 3.2 Biokinetic model parameter estimation and experimental validation

The biokinetic model parameters (**Table S2**) were estimated by minimizing the root mean square error (RMSE) between predicted and experimental time-resolved metabolite concentrations. These parameters include the maximum biomass uptake rate for acid production, substrate affinity constants for biomass, maximum specific production rate of alcohols, substrate saturation constants, biomass decay rate, and yield coefficients. Due to the limitations of small-scale systems (e.g., serum bottles), where limited sample volume restricts the simultaneous measurement of substrate concentrations in both gas and dissolved phases, only metabolite profiles in liquid fermentation were used for model calibration. Initial parameter estimates for chain elongation were derived from literature studies on C_2_–C_6_ acid production from CO_2_/H_2_ mixtures in membrane biofilm reactors (Chen and Ni, 2016).

The time-resolved metabolite concentration profiles obtained from 0D dynamic-kinetic model simulations, using the calibrated model parameters, were presented in **Fig. 3**. The simulated profiles exhibit good alignment with the experimental metabolite production over time, capturing key phases of gas fermentation, including biomass growth and decay, as well as the acetogenesis and solventogenesis processes. The RMSE value of 0.18 which falls within the acceptable range of <0.30 suggests reasonable accuracy of the model to predict experimental trends. Notably, the model successfully reproduced the maximum biomass concentration (0.25 g/L at ∼24 hours) and the final yields of the C_2_–C_6_ acids and alcohols at 120 hours. This agreement suggests that the kinetic framework can capture the microbial metabolic shifts under CO_2_/H_2_ conditions. The ability to simulate both acidogenesis and solventogenesis further supports previous studies indicating that acetogens regulate electron flow to optimize carbon flux, with H_2_ availability influencing the distribution of reduced products (Hermann et al., 2020; Zhu et al., 2020). Moreover, the model’s predictive capability highlights the robustness of parameter estimation, ensuring reliable representation of the gas fermentation process across varying operating conditions. This is particularly relevant for process scale-up, where accurate kinetic modelling is essential for optimizing bioreactor performance and product selectivity (Singh et al., 2024).

**Fig. 3.**
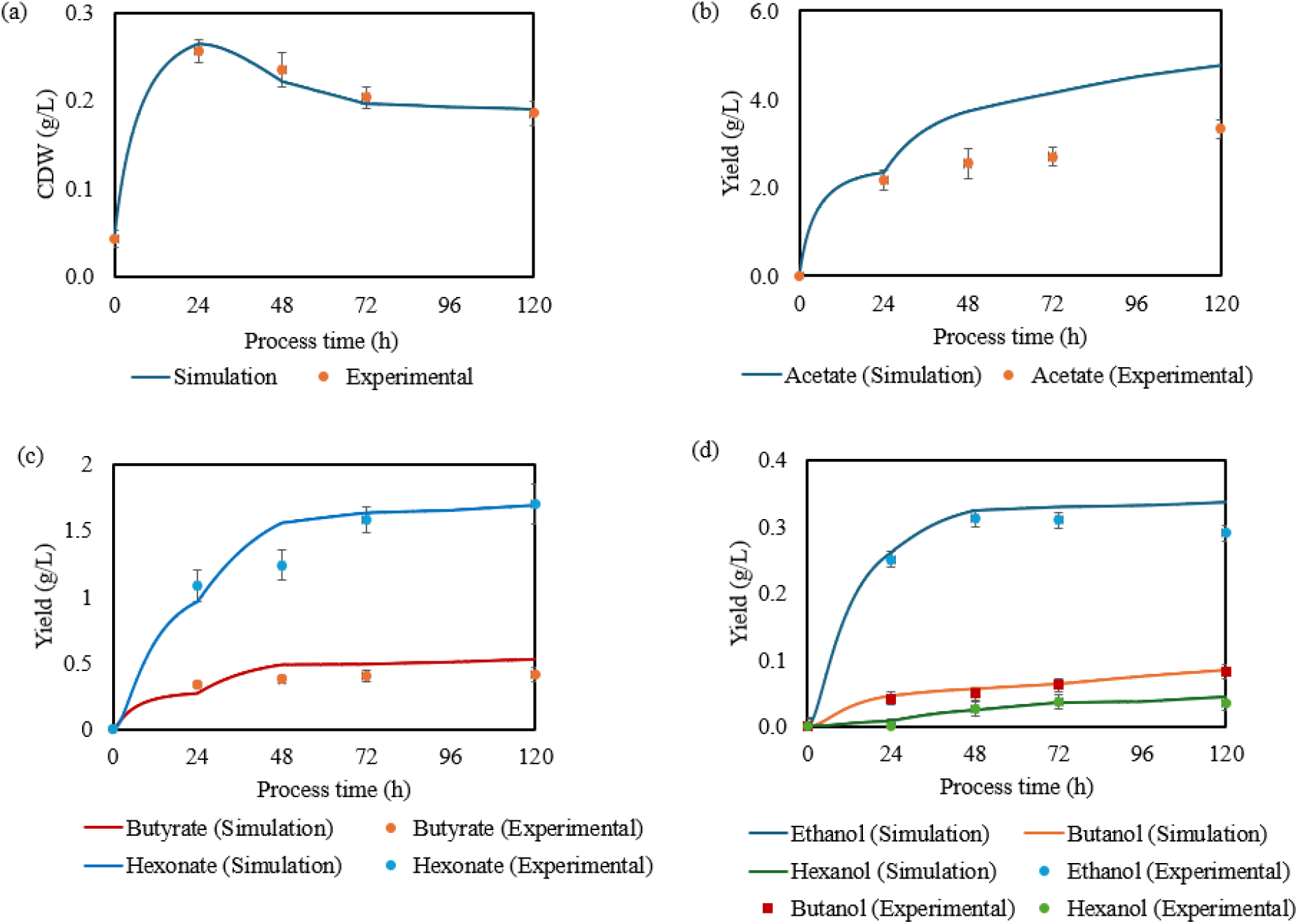
Model predicted time-resolved profiles of bioconversion products along with experimental data: **(a)** Cell growth; Production yields of **(b)** acetate, **(c)** butyrate and hexanoate, as well as **(d)** ethanol, butanol and hexanol.

### 3.3 Effect of the liquid-to-gas volume ratio (V_L_/V_G_) on metabolite formation

The computational model was further applied to investigate the effect of the liquid-to-gas volume ratio (V_L_/V_G_) on metabolite formation, providing quantitative insights into biomass growth, dissolved H_2_ concentration and metabolite distribution. Simulated total acid and alcohol concentrations at 24, 48, and 72 hours revealed that increasing the liquid volume enhances metabolite production, as presented in **Fig. 4a**. These model predictions were further validated by experimental fermentation under CO_2_/H_2_ gas conditions (**Fig. 4b**). Biomass growth followed a similar trend to product formation (**Fig. S3a**), increasing with higher V_L_/V_G_, suggesting that larger liquid volumes improve gas solubility and substrate availability, thereby enhancing microbial metabolism. The larger liquid phase improves mass transfer rates by increasing the contact area between gas and liquid, reducing diffusion limitations (Lanzillo et al., 2020). The dissolved H_2_ concentration profiles showed a gradual decline over time, with the lowest concentration at V_L_/V_G_ = 0.25 and the highest at V_L_/V_G_ = 4 (**Fig. S3b**). This trend emphasize the importance of liquid volume in stabilizing gas uptake rates, as a larger liquid phase can buffer against rapid depletion of dissolved H_2_ by acting as a reservoir, thereby sustaining microbial metabolism for extended periods (Rahimi et al., 2018). Conversely, in lower V_L_/V_G_ conditions, H_2_ diffusion may be insufficient to match microbial consumption rates, leading to premature depletion and suboptimal metabolic activity (Ruggiero et al., 2022). These findings highlight the importance of gas-liquid mass transfer, where a larger liquid phase facilitates better substrate diffusion, ultimately boosting metabolic efficiency.

**Fig. 4.**
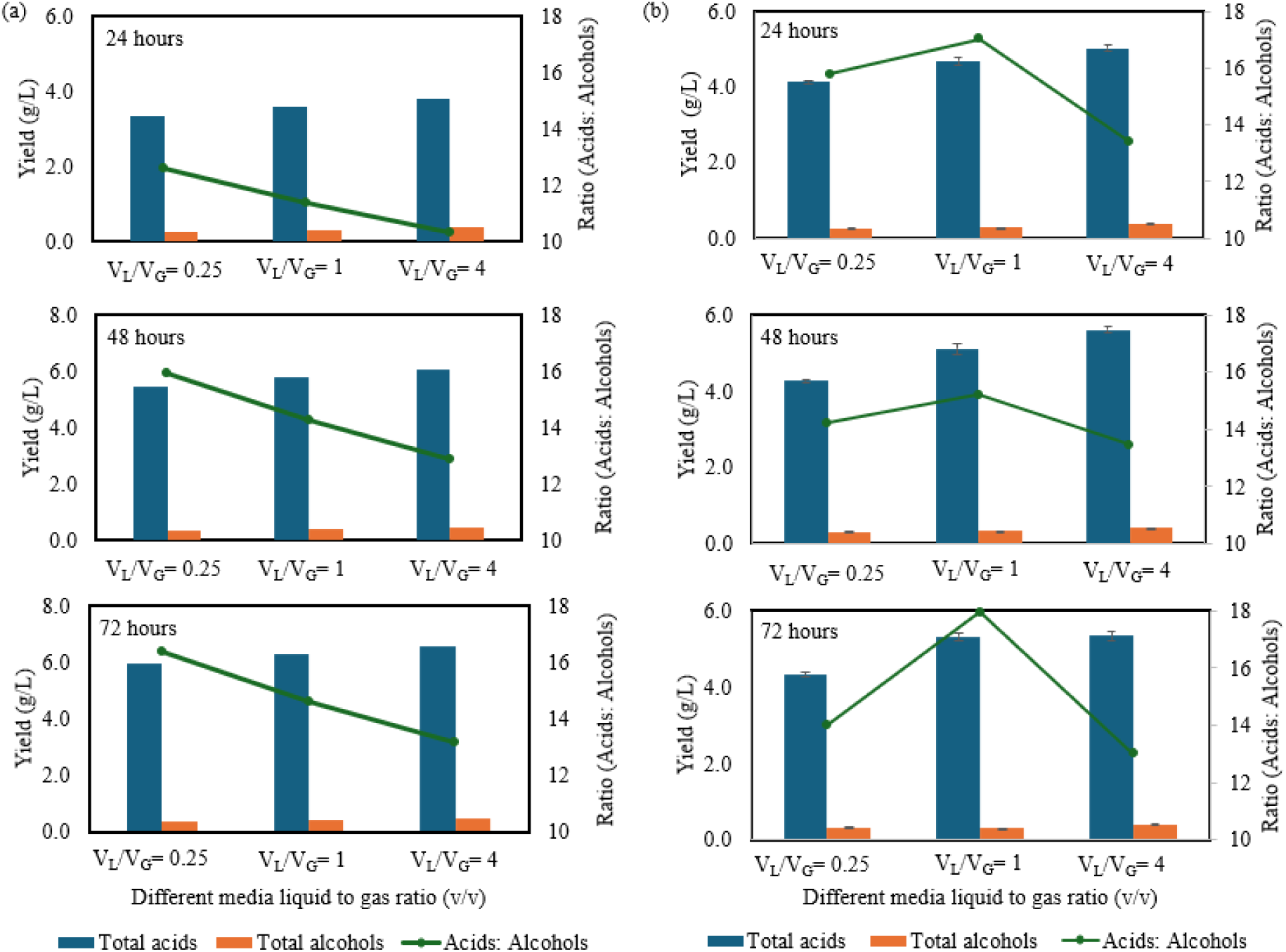
Total acid and alcohol yields, and the corresponding acid-to-alcohol ratios at 24, 48, and 72 hours based on **(a)** Simulation, and **(b)** Experimental results from *C. carboxidivorans* sp. 624 fermentation using CO_2_/H_2_ (20/80) in RCM medium at 25 °C.

Unsurprisingly, the total acids and alcohols, as well as the acid-to-alcohol ratio, were strongly influenced by the H_2_/CO_2_ feedstock ratio (80:20) and the WLP (**Fig. 4b**). At 72 hours, V_L_/V_G_ = 4 conditions yielded the highest total acids (5.34 g/L) and alcohols (0.41 g/L), highlighting the role of an H_2_-rich environment in enhancing acetogenesis and chain elongation. The slightly lower acid-to-alcohol ratio (13) at V_L_/V_G_ = 4 conditions indicates higher acid conversion to alcohols, supporting the hypothesis of product diversification through excess hydrogenation. Conversely, at V_L_/V_G_ = 1, the fermentation produced the highest acid-to-alcohol ratio (18), favouring acetate production as moderate H_2_ availability promoted acetogenesis while minimizing solventogenesis. Collectively, these findings demonstrate the interplay between gas-liquid interactions, substrate availability, and metabolic pathways in CO_2_/H_2_ fermentation. The computational model provides predictive insights that align with experimental observations, reinforcing the significance of optimizing V_L_/V_G_ and electron donor-to-acceptor ratios to tailor product distribution in CO_2_-based bioprocesses.

### 3.4 CFD simulations and validation

As we observed the impact of gas-liquid interactions, the importance of spatial distribution becomes more pronounced. A detailed understanding of the non-uniform spatial distributions of key components, such as substrates and metabolites, is essential, as these factors significantly impact cell growth and final bioproduction yields. To account for both temporal variations and spatial distributions within the bioreactor, the biokinetic model’s estimated parameters were integrated with computational fluid dynamics (CFD) simulations. In small-scale batch systems, like serum bottles agitated at 200 rpm in this study, spatial variations and convective mixing effects are predicted to be minimal (Zhang et al., 2010). This results in a largely good qualitative and quantitative agreement between 0D dynamic-kinetic model estimations and CFD simulations (**Fig. 5**), where time-resolved concentration profiles of biomass and metabolites are compared.

**Fig. 5.**
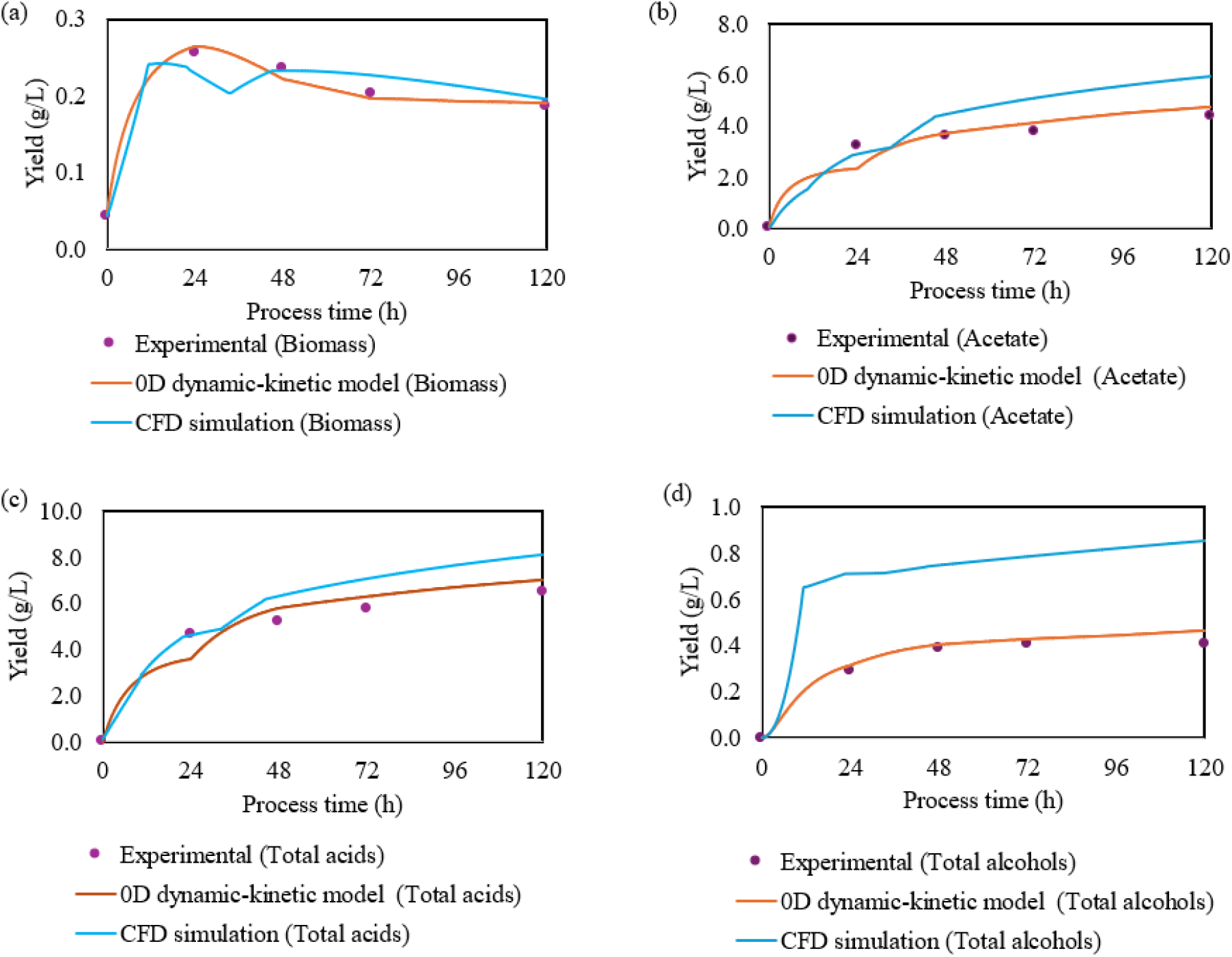
Profiles calculated from 0D dynamic-kinetic model and integrated CFD simulations compared with experimental data (as in **Fig. 3**), **(a)** Cell growth (biomass), **(b)** Acetate, **(c)** Total acids, and **(d)** Total alcohols.

The concentration contours of biomass and acetate from 3D CFD simulations of the serum bottle model were analysed at different time points (5, 18, and 72 hours) during the *Clostridium carboxidivorans* sp. 624 fermentation under CO_2_/H_2_ (20/80) in RCM media (**Fig. 6a**). Simulations indicated a rise in biomass concentration from 5 to 18 hours (**Fig. 6b**).

**Fig. 6.**
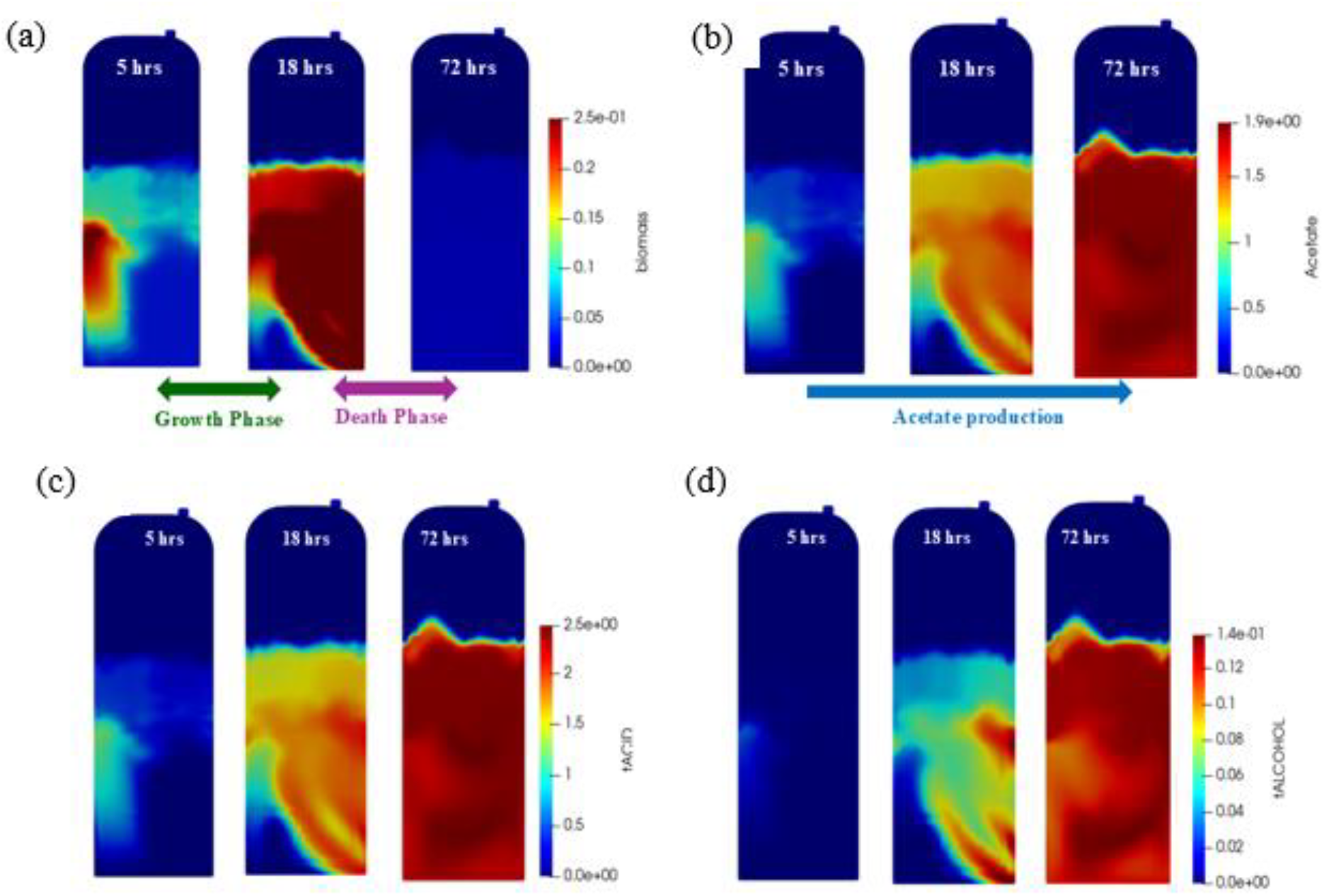
Concentration contours at different time points of reaction **(a)** Cell growth (biomass), and **(b)** Acetate, **(c)** Total acids, and **(d)** Total alcohols.

Although acetate is later converted into ethanol, its sustained accumulation suggests its role as a key intermediate in the fermentation process. The spatial distribution patterns of biomass and acetate closely mirrored each other, indicating that acetate biosynthesis is linked to microbial growth, with active metabolism facilitating continuous acetate generation (Kurt et al., 2023).

At 5 hours, acids production dominates while alcohol levels are negligible, indicating the early acetogenic phase in the time-resolved concentration profiles (**Fig. 6c**). By 18 hours, acid production significantly exceeds that of alcohols, confirming the predominance of acetogenesis over solventogenesis. At 72 hours, both acids and alcohols production reach their maximum level, reflecting the transition to solventogenesis as more reducing equivalents become available for alcohol formation (**Fig. 6d**). The temporal distinction between metabolic phases corroborates with previous studies on syngas fermentation, where an extended acetogenic phase precedes solventogenesis under excess H_2_ conditions (Lanzillo et al., 2020).

Overall, these results highlight the potential of CFD simulations to support gas fermentation process development. Factors such as spatial heterogeneities in substrate distribution, gas-liquid mass transfer, and hydrodynamics that crucially influence bioprocess efficiency can be captured in the model. It is worth acknowledging that application of serum bottle scale model may be limited as larger scale continuous stirred-tank reactors (CSTRs) undergo higher stirring frequencies and amplitudes. In CSTRs, physical and mechanical non-uniformities are more evident which could lead to greater discrepancies between CFD predictions and traditional 0D first-principles approaches (Hanspal et al., 2023). In larger bioreactors, factors such as mixing intensity, bubble dynamics, and localized nutrient gradients can lead to non-uniform microbial activity, potentially limiting productivity (Singh et al., 2024). Future research will expand therefore these simulations to pilot-scale systems to gain deeper insights into fluid flow behaviour, optimize gas-to-liquid transfer efficiency, and refine operational strategies for improved metabolite yields in industrial applications.

## 4. Conclusion

This study demonstrates the potential of *Clostridium carboxidivorans* sp. 624 for CO_2_/H_2_ assimilation using a hybrid experimental and *in silico* and approach. A significantly higher acetate (3.33 g/L) and hexanoate (1.70 g/L) yields under CO_2_/H_2_ in RCM conditions, relative to the N_2_ control, alludes to the role of gas-phase substrates in driving acetogenesis. The kinetic model and experimental validation highlight the crucial importance of the high liquid-to-gas volume ratio (V_L_/V_G_) and gas interactions in enhancing metabolite production and microbial metabolism during CO_2_/H_2_ fermentation. Integrated kinetic modelling and 3D CFD simulations also informed bioprocess design for advancing scalable, carbon-negative CO_2_ biomanufacturing.

## Acknowledgements

The authors gratefully acknowledge financial support from Agency for Science, Technology and Research (A*STAR), Singapore (C233017002 and Singapore Integrative Biosystems and Engineering Research Strategic Research & Translational Thrust (SIBER SRTT). The computational work for this article was partially performed on resources of the National Supercomputing Centre (NSCC), Singapore (https://www.nscc.sg).

## Notes

### Competing Interest Statement

The authors have declared no competing interest.

### Summary of Updates

Abstract updated, manuscript updated, supplemental files updated.

