## Supplementary material for "An Experimental and Multiphysics Simulations Study of *Clostridium carboxidivorans* sp. 624 for Acid and Alcohol Production in CO_2_/H_2_": SI

### Figure S1

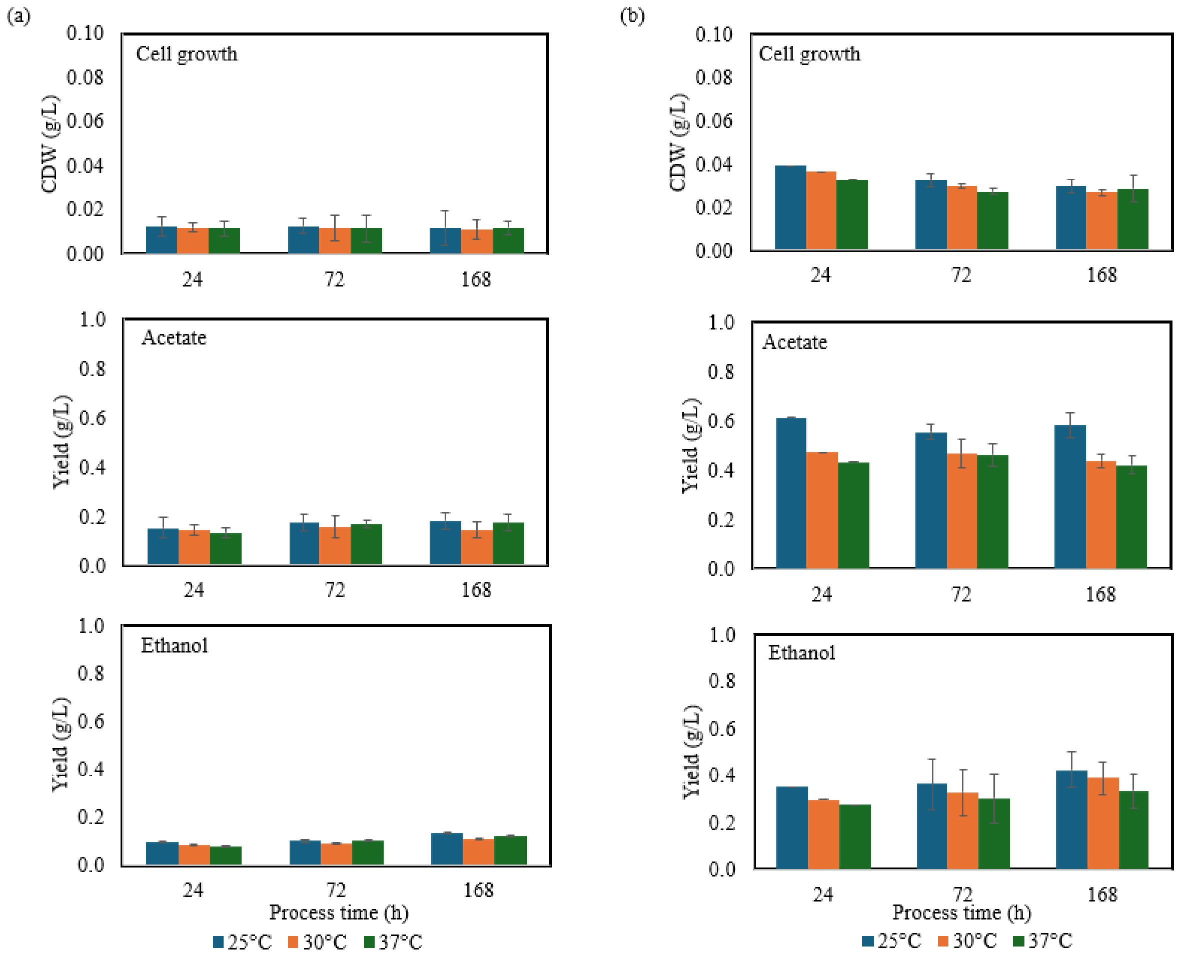


**Fig. S1.** Cell growth and production of $C_2$ metabolite analysis of *C. carboxidivorans* sp. 624 fermentation under **(a)** $N_2$, and **(b)** $CO_2/H_2$ (20/80) in PETC media at 25°C, 30°C, and 37°C.

**Figure S2**

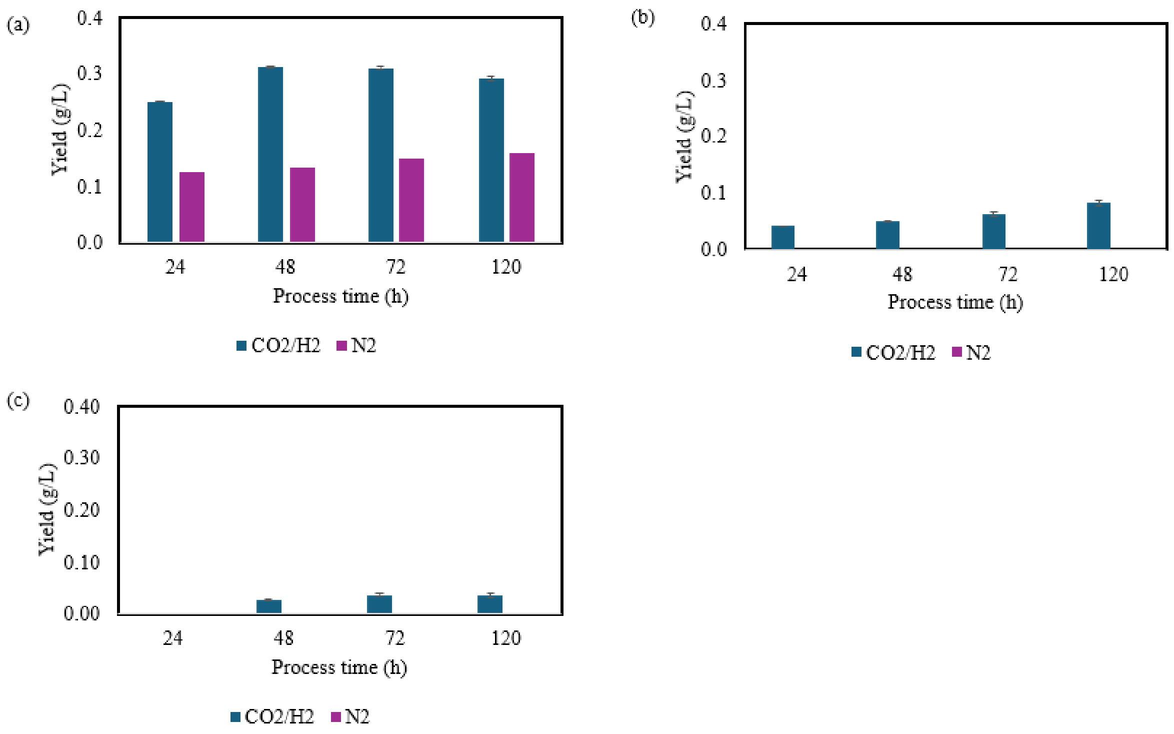


**Fig. S2.** *C. carboxidivorans* sp. 624 fermentation with $CO_2/H_2$ in RCM medium at 25°C: Production yields of **(a)** Ethanol, **(b)** Butanol, and **(c)** Hexanol.

**Figure S3**

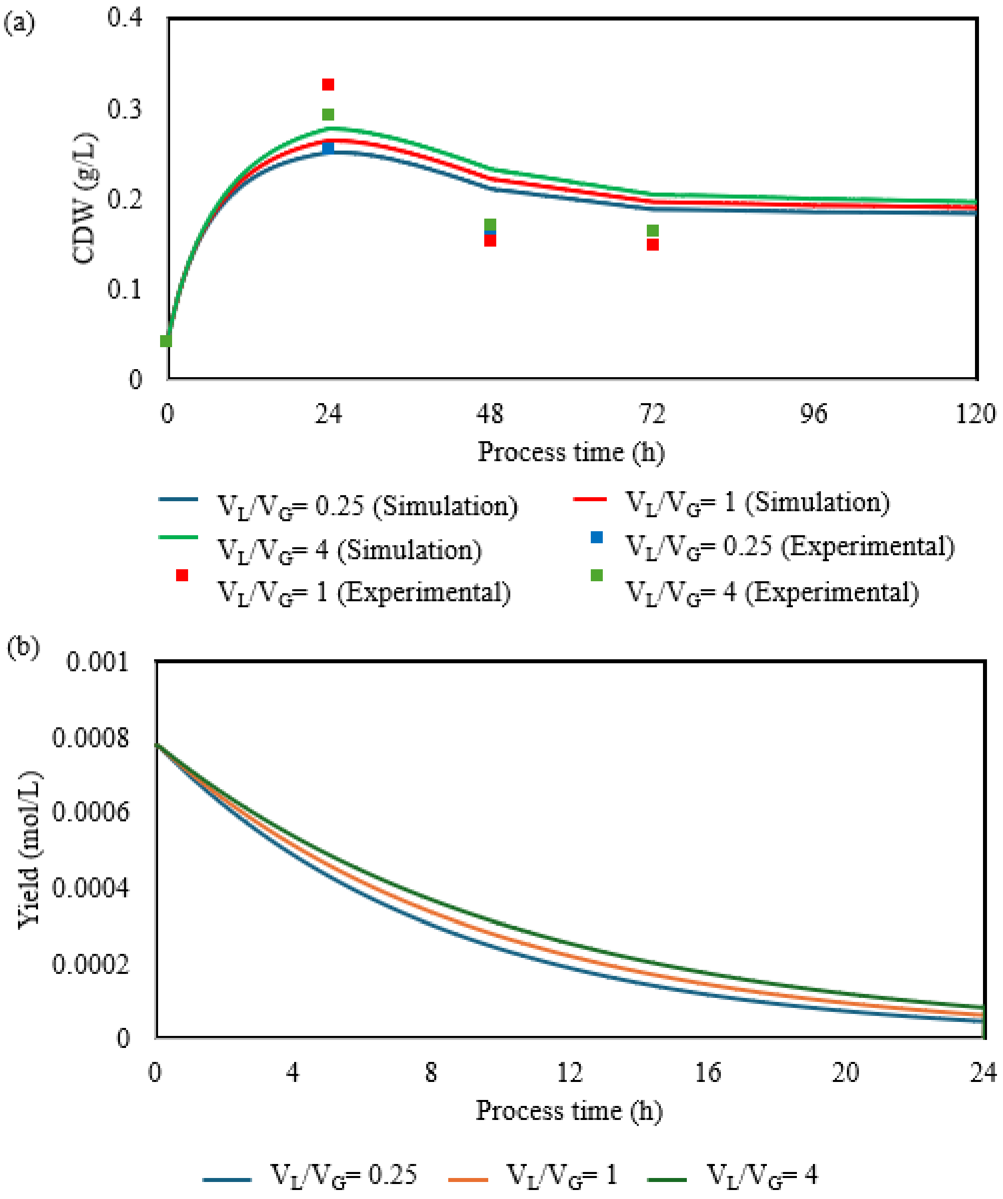


**Fig. S3.** Time resolved profiles of **(a)** biomass and **(b)** equilibrated $H_2$ amount in liquid media at different $V_L/V_G$.

**Table S1.**

First-principles-based mass balance equations for gas and liquid phase components.

| | |
|---|---|
| $V_G \frac{dP_{H2}}{dt} = -V_L RT(k_L a)_{H2}(P_{H2}KH_{H2} - C_{H2L})$ | $(S1.1)$ |
| $\frac{dC_{H2L}}{dt} = RT(k_L a)_{H2}(P_{H2}KH_{H2} - C_{H2L}) + r_{H2}$ | $(S1.2)$ |
| $\frac{dC_X}{dt} = r_X = (q_{AC}Y_{X/AC} + q_{BA}Y_{X/BA} + q_{HA}Y_{X/HA} - r_d)C_X$ | $(S1.3)$ |
| $\frac{dC_{AC}}{dt} = r_{AC} = (q_{AC} - q_{BA}Y_{AC/BA} - q_E Y_{AC/E})C_X$ | $(S1.4)$ |
| $\frac{dC_{BA}}{dt} = r_{BA} = (q_{BA} - q_{HA}Y_{BA/HA} - q_B Y_{BA/B})C_X$ | $(S1.5)$ |
| $\frac{dC_{HA}}{dt} = r_{HA} = (q_{HA} - q_H Y_{HA/H})C_X$ | $(S1.6)$ |
| $\frac{dC_E}{dt} = r_E = q_E C_X$ | $(S1.7)$ |
| $\frac{dC_B}{dt} = r_B = q_B C_X$ | $(S1.8)$ |
| $\frac{dC_H}{dt} = r_H = q_H C_X$ | $(S1.9)$ |

**Table S2.**

Calibrated biokinetic model parameters.

| Parameter | Description of the parameter | Value | Unit |
|---|---|---|---|
| | Kinetic Parameters | | |
| $\boldsymbol{\mu_{Ac}}$ | Maximum uptake rate of biomass for acetate production | 9.5 | 1/h |
| $\boldsymbol{\mu_{BA}}$ | Maximum uptake rate of biomass for butyrate production | 0.778 | 1/h |
| $\boldsymbol{\mu_{HA}}$ | Maximum uptake rate of biomass for butyrate production | 2.625 | 1/h |
| $\boldsymbol{K_{H2}^{i}}$ $(\boldsymbol{i} = \boldsymbol{AC}, \boldsymbol{BC}, \boldsymbol{HA})$ | $H_2$ affinity constant for biomass for *i* production | 1.16 $e^{-6}$ | $molL^{-1}$ |
| $\boldsymbol{k_{Ac}}$ | Acetate affinity constant for biomass | 0.0094 | $molL^{-1}$ |
| $\boldsymbol{k_{BA}}$ | Butyrate affinity constant for biomass | 0.0022 | $molL^{-1}$ |
| $\boldsymbol{\mu_{E}}$ | Maximum production rate of ethanol | 9.6 | 1/h |
| $\boldsymbol{\mu_{B}}$ | Maximum production rate of butanol | 6.4 | 1/h |
| $\boldsymbol{\mu_{H}}$ | Maximum production rate of hexanol | 0.4 | 1/h |
| $\boldsymbol{K_{H2}^{j}}$ $(\boldsymbol{i} = \boldsymbol{E}, \boldsymbol{B}, \boldsymbol{H})$ | Saturation constant for $H_2$ for *j* production | 2.2 $e^{-4}$ | $molL^{-1}$ |
| $\boldsymbol{k_{UAc}}$ | Saturation constant for acetate | 0.00075 | $molL^{-1}$ |
| $\boldsymbol{k_{UBA}}$ | Saturation constant for butyrate | 0.0005 | $molL^{-1}$ |
| $\boldsymbol{k_{UHA}}$ | Saturation constant for hexanoate | 0.0005 | $molL^{-1}$ |
| $\boldsymbol{k_{dec}}$ | Decay rate coefficient for biomass | 0.1294 | 1/h |
| | Yield | | |
| Y(X/AA) | yield coefficient of biomass with respect to acetate | 0.1486 | |
| Y(X/BA) | yield coefficient of biomass with respect to butyrate | 0.0891 | |
| Y(X/HA) | yield coefficient of biomass with respect to hexanoate | 0.0557 | |

| | | |
|---|---|---|
| Y(AA/BA) | yield coefficient of acetate with respect to butyrate | 0.4 |
| Y(BA/HA) | yield coefficient of butyrate with respect to hexanoate | 0.625 |
| Y($H_2$/AA) | yield coefficient of $H_2$ with respect to acetate | 0.5743 |
| Y($H_2$/BA) | yield coefficient of $H_2$ with respect to butyrate | 0.3445 |
| Y($H_2$/HA) | yield coefficient of $H_2$ with respect to hexanoate | 0.2154 |
| Y($H_2$/E) | yield coefficient of $H_2$ with respect to ethanol | 2 |
| Y($H_2$/B) | yield coefficient of $H_2$ with respect to butanol | 2 |
| Y($H_2$/H) | yield coefficient of $H_2$ with respect to hexanol | 4 |
| Y(AA/E) | yield coefficient of acetate with respect to ethanol | 1 |
| Y(BA/B) | yield coefficient of butyrate with respect to butanol | 1 |
| Y(HA/H) | yield coefficient of hexanoate with respect to hexanol | 1 |
